# Trafficking Machinery is Rapidly Primed to Facilitate Polarised IL-6 Secretion in Dendritic Cells

**DOI:** 10.1101/2023.07.13.548819

**Authors:** Harry Warner, Tongxiang Chen, Shweta Mahajan, Martin ter Beest, Peter Linders, Giulia Franciosa, Frans Bianchi, Geert van den Bogaart

**Affiliations:** Department of Molecular Immunology, Groningen Biomolecular Sciences and Biotechnology Institute, University of Groningen, The Netherlands; Division of Immunobiology, Center for Inflammation and Tolerance, Cincinnati Children’s Hospital, Cincinnati, Ohio, USA; Department of Tumour Immunology, Radboud Institute for Molecular Life Sciences, Radboud University Medical Center, The Netherlands; Novo Nordisk Foundation Center for Protein Research, Faculty of Health and Medical Sciences, Copenhagen University, Copenhagen, Denmark

## Abstract

The mounting of an adaptive immune response is critical for removing pathogens from the body and generating immunological memory. Central to this process are myeloid cells, which sense pathogens through a variety of cell surface receptors, engulf and destroy pathogens and become activated. Activation is essential for the release of cytokines as well as the cell-surface presentation of pathogen-derived-antigens. Activation-induced cytokine release by myeloid cells requires a complex series of molecular events to facilitate cytokine expression. However, although the transcriptional machinery regulating cytokine expression is well defined, it is becoming increasingly clear that trafficking machinery has to be re-programmed through post-translational modifications to dynamically regulate cytokine secretory events. We demonstrate through quantitative total internal-resonance fluorescence (TIRF) microscopy that short-term stimulation with the pathogenic stimulus lipopolysaccharide (LPS) is sufficient to up-regulate IL-6 secretion rates in human blood monocyte-derived dendritic cells and that this secretion is asymmetric and thus polarised. Using bioinformatics analysis of our phosphoproteomic data, we demonstrate that LPS stimulation of monocyte-derived dendritic cells rapidly reprograms SNARE-associated membrane trafficking machinery, through phosphorylation/dephosphorylation events. Finally, we link this enhanced rate of secretion to the phosphorylation of the SNARE protein VAMP3 at serine 44 (48 in mice), by showing that this phosphorylation drives the release of VAMP3 by its chaperone WDFY2 and the complexing of VAMP3 with STX4 at the plasma membrane.

## Introduction

The activation of myeloid cells in response to a pathogenic insult requires a complex series of molecular events for choreographing a local response to a pathogenic invasion and the generation of an adaptive immune response by B and T cells (Patente et al., 2019). In order to coordinate these events, myeloid cells release soluble signals in the form of cytokines to surrounding tissue at the site of infection and at immunological synapses formed with T cells. Whilst elegant genetic studies have identified key cell-surface receptors and transcription factors that drive the expression of cytokines, less is known about the short-term response of myeloid cells to pathogenic insults. Furthermore, although it is known that trafficking machinery is utilised to facilitate cytokine secretion, it has long been thought that cytokines utilise pre-established membrane trafficking circuits, the same as utilised by inactive myeloid cells (Revelo et al., 2019).

Soluble N-ethylmaleimide-sensitive factor attachment protein receptors (SNARE) proteins are key regulators of membrane traffic, that confer both identity to their associated membrane and drive membrane fusion(Jahn and Scheller, 2006)(Hong, 2005)(Koike and Jahn, 2022). Both of these functions are achieved through the specific complexing of Q-SNAREs with a cognate R-SNARE, which drives fusion of the associated membranes. Whilst this function of SNAREs has long been appreciated, it’s becoming increasingly apparent that this Q-SNARE-R-SNARE specificity is plastic, and can be rapidly modified through post-translational modifications(Warner, Mahajan and van den Bogaart, 2022). These modifications can in turn be coordinated in response to an ever-changing extracellular environment.

Cell polarity is a key means to couple the internal architecture of a cell to the cell’s external environment. Different polarity networks enable cells to effectively perform their functions. For instance, epithelial cell plasma membranes are separated into apical and basolateral domains, whereas fibroblasts and myeloid cells establish front-rear polarity. Membrane traffic has long been known to support these different cellular polarities(Murray, Kay, Sangermani and Stow, 2005)(Mostov, Su and ter Beest, 2003), ensuring the correct positioning of transmembrane proteins within the plasma membrane and ensuring that key extracellular proteins are secreted at the correct location.

We have previously shown that the trafficking machinery plays an active role in controlling interleukin (IL)-6 synthesis; a pool of IL-6 at the perinuclear recycling compartment signals via its receptor IL-6R to the genome to dampen IL-6 transcription(Verboogen et al., 2019). Thus, the trafficking machinery can feature as part of a larger feedback loop to fine-tune extracellular cytokine concentrations. This is likely critical, as IL-6 is a key component in the initiation of sepsis (Hack et al., 1989), promotes survival of chromosomally instable cancers(Hong, Schubert, Tijhuis, Requesens, Roorda, van den Brink, Ruiz, Bakker, van der Sluis, Pieters, Chen, Wardenaar, van der Vegt, Spierings, de Bruyn, van Vugt and Foijer, 2022), and is a biomarker for the progression of metastatic breast cancer (Salgado et al., 2003) and COVID19 – associated hyperinflammation (Santa Cruz et al., 2021). Furthermore, VAMP3, an R-SNARE protein regulating trafficking of recycling endosomes (REs), is not only essential for IL-6 secretion (Boddul et al., 2014; Manderson et al., 2007; Verboogen et al., 2019), but is also implicated as a driver of inflammatory bowel disease (Ventham et al., 2013).

We also previously observed increased complexing between VAMP3 and the Q-SNARE STX4 at the plasma membrane in dendritic cells, in response to stimulation with the pathogenic stimulus lipopolysaccharide (LPS) (Verboogen et al., 2017). However, the extent to which the trafficking machinery is re-programmed to facilitate cytokine secretion is still unclear. Furthermore, previous work on cytokine secretion has suggested that IL-6 secretion is unpolarised in myeloid cells(Manderson et al., 2007). However, VAMP3 is typically deployed by cells to achieve polarised delivery of trafficking cargo to set membrane domains(Fields et al., 2007). Furthermore, IL-6 secretion by epithelial cells has been reported as polarised towards the apical membrane (Healy et al., 2015). Therefore, it is unclear if VAMP3 is utilised by dendritic cells to achieve non-polarised IL-6 secretion. However, polarised secretion would be logical, as this may for instance facilitate secretion of IL-6 at immunological synapses with T cells, as IL-6 is a T cell survival and polarisation factor (Li et al., 2018).

Here, we utilise total internal-resonance fluorescence (TIRF) microscopy to address these questions. We demonstrate that whilst IL-6 positive vesicles are evenly distributed throughout dendritic cells, their secretion pattern is asymmetric and thus partially polarised. Furthermore, we show that brief LPS stimulation (1-4 hours) is sufficient to increase the rate of IL-6 secretion, indicating that trafficking machinery is rapidly post-translationally modified to promote cytokine secretion. Finally, we link this up-regulated IL-6 traffic to the phosphorylation of VAMP3 at serine 44 (48 in mice), which promotes complexing of VAMP3 with STX4 at the plasma membrane, ultimately driving IL-6 secretion.

## RESULTS

### TIRF Microscopy Reveals Spatio-temporal Dynamics of IL-6 Secretion

To differentiate between short-term and long-term changes to trafficking machinery, we overexpressed GFP-tagged IL-6 in human peripheral blood monocyte-derived dendritic cells and imaged secretory events in the absence or short-term presence of LPS using total internal reflection fluorescence (TIRF) microscopy. GFP fluorescence is partially quenched within the relatively acidic environment of vesicles; however this quenching is relieved upon exocytosis, creating a spark of fluorescence (Miesenböck et al., 1998). These sparks can then be visualized using TIRF, as has been previously described for GFP-tagged IL-6 (Verboogen et al., 2018). This revealed that short-term stimulation (imaging between 1 – 4 hours after LPS addition) of dendritic cells was sufficient to up-regulate the rate of IL-6 secretion (Fig 1A and 1B). This time frame was selected as we previously observed that LPS induced IL-6 accumulates in the recycling endosome prior to secretion, with accumulation peaking at 4 hours (Verboogen et al, 2019).

**Figure 1.**
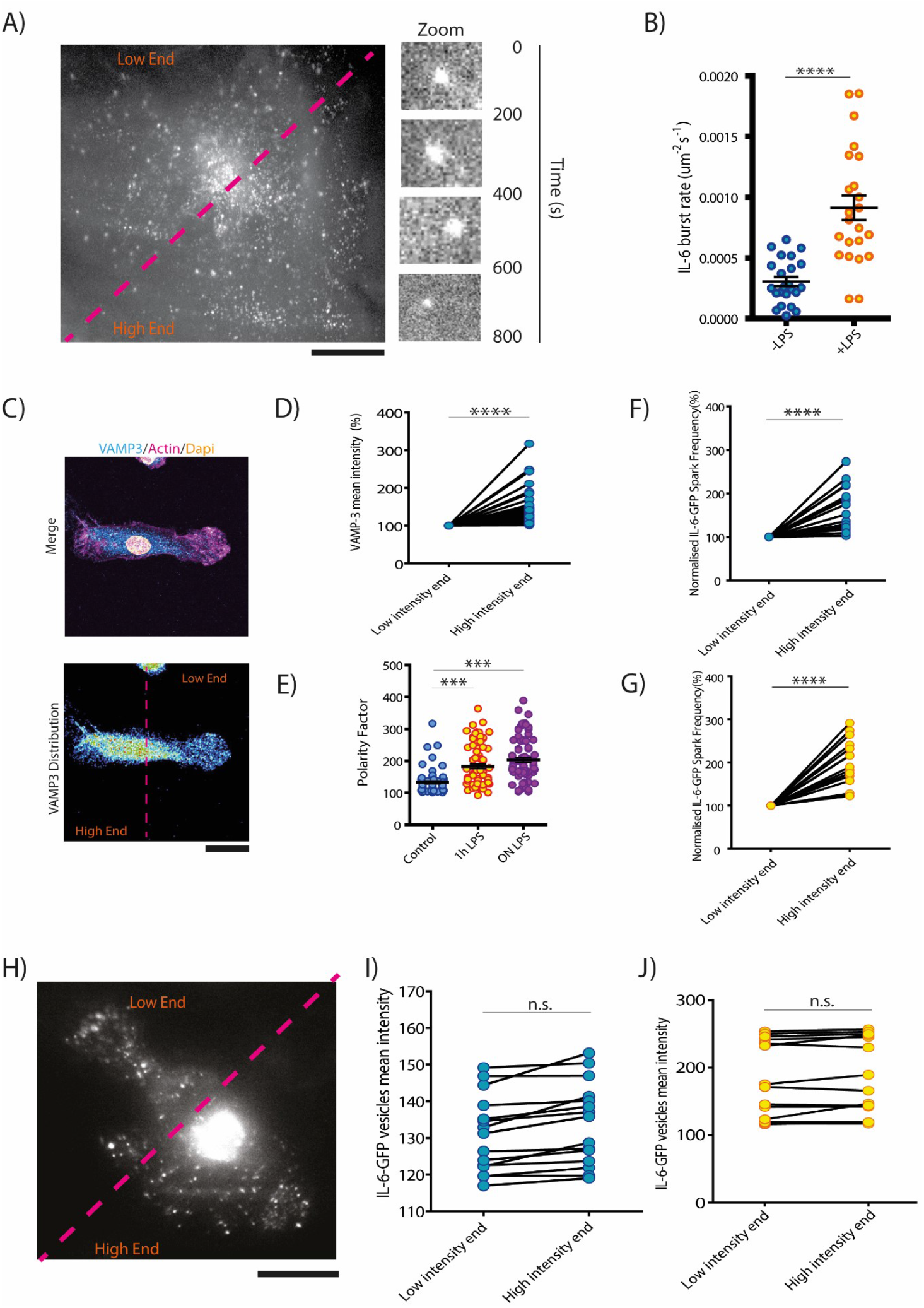
IL-6 Trafficking Apparatus is Polarised and Regulated by LPS Stimulation. A) Example TIRF imaging of IL-6 secretory events. Secretory sites are polarised along indicated purple line (arbitrary line segmenting the cell). B) Quantification of IL-6 secretion rate in the absence or presence of short-term LPS stimulation (23 measurements per condition over 3 donors). C) Example image of a dendritic cell immunostained for VAMP3. D) Quantification of VAMP3 staining at high and low ends of the cell display confirms that VAMP3 is polarised in its distribution (67 measurements over 3 donors). E) VAMP3 polarisation increases after 1 hour and overnight LPS stimulation (minimum of 67 measurements condition per over 3 donors). F) Quantification of IL-6 secretion rate at “high” and “low” ends of the cell in inactive dendritic cells. (19 measurements over 3 donors) G) Quantification of IL-6 secretion rate at “high” and “low” ends of the cell in activated dendritic cells (17 measurements over 3 donors). H) Example TIRF imaging of IL-6 positive vesicles. I) Quantification of IL-6+ vesicles at “high” and “low” ends of unstimulated dendritic cells (15 measurements over 3 donors). J) Quantification of IL-6+ vesicles at “high” and “low:” ends of dendritic cells stimulated with LPS for ∼ 1-4 hours prior to imaging (14 measurements over 3 donors; Scale bars indicate 20 microns). Statistical significance calculated using a 2-sided unpaired t-test for panels B, D, F, G, I and J; and ANOVA/Tukey multiple comparison test for panel E. ***, P < 0.001; ****, P<0.0001; n.s., not significant.

In order to understand the spatio-temporal dynamics of IL-6 secretion we first confirmed that VAMP3 (required for IL-6 secretion) is polarised in myeloid cells, as has been previously observed for macrophages (Veale et al., 2011). Arbitrary segmenting of confocal micrographs of macrophages stained for VAMP3, revealed an asymmetric, polarised distribution of VAMP3+ vesicles. (Fig. 1C – 1E). We therefore calculated the polarity index of VAMP3+ vesicle distribution (high-end intensity/low-end intensity X100). We also performed a similar sub-cellular analysis to determine if the distribution of IL-6 secretion sites is polarised. This revealed that IL-6 secretion is asymmetric, which would allow a dendritic cell *in vivo* to coordinate secretory events with its local environment (Fig. 1F and 1G). However, this conflates with previous work that showed that IL-6 secretion is unpolarised, in contrast to TNF-α secretion, which is polarised to the phagocytic cup (Manderson et al., 2007). We therefore checked the distribution of IL-6+ vesicles within the cytosol, using TIRF microscopy (Fig 1I-J). This revealed that the distribution of intracellular IL-6 is unpolarised. Thus, although IL-6 distribution is non-polarised, the final secretory event is.

### LPS Stimulation Primes Trafficking Machinery Via VAMP3 Phosphorylation

Having established that short-term LPS stimulation can drive IL-6 secretion, we examined our human dendritic cell phosphoproteomic data (Warner *et al*., 2023) to identify novel phosphorylation/dephosphorylation events that regulate SNAREs and SNARE associated proteins, using STRING software (Szklarczyk et al., 2019). This revealed a network of proteins that interact with VAMP3 (including VAMP3 itself) that are rapidly phosphorylated in response to LPS stimulation (Fig. 2A). Further inspection of the data set revealed two VAMP3 phosphorylation events. One at serine 44 (significant at 1 and 4 hours) (Fig 2B) and the other at serine 11 (significant at 4 hours) (Fig 2C). Furthermore, consistent with our previous work (Verboogen et al., 2017), we could not detect a change in VAMP3 protein levels (Fig 2D) in our total cell proteomics data. We therefore chose to follow up the serine 44 phosphorylation event as not only is this phosphorylation more significant, but it is also more rapid (Fig 2E). These findings suggest that VAMP3 phosphorylation might prime dendritic cells for IL-6 secretion.

**Figure 2.**
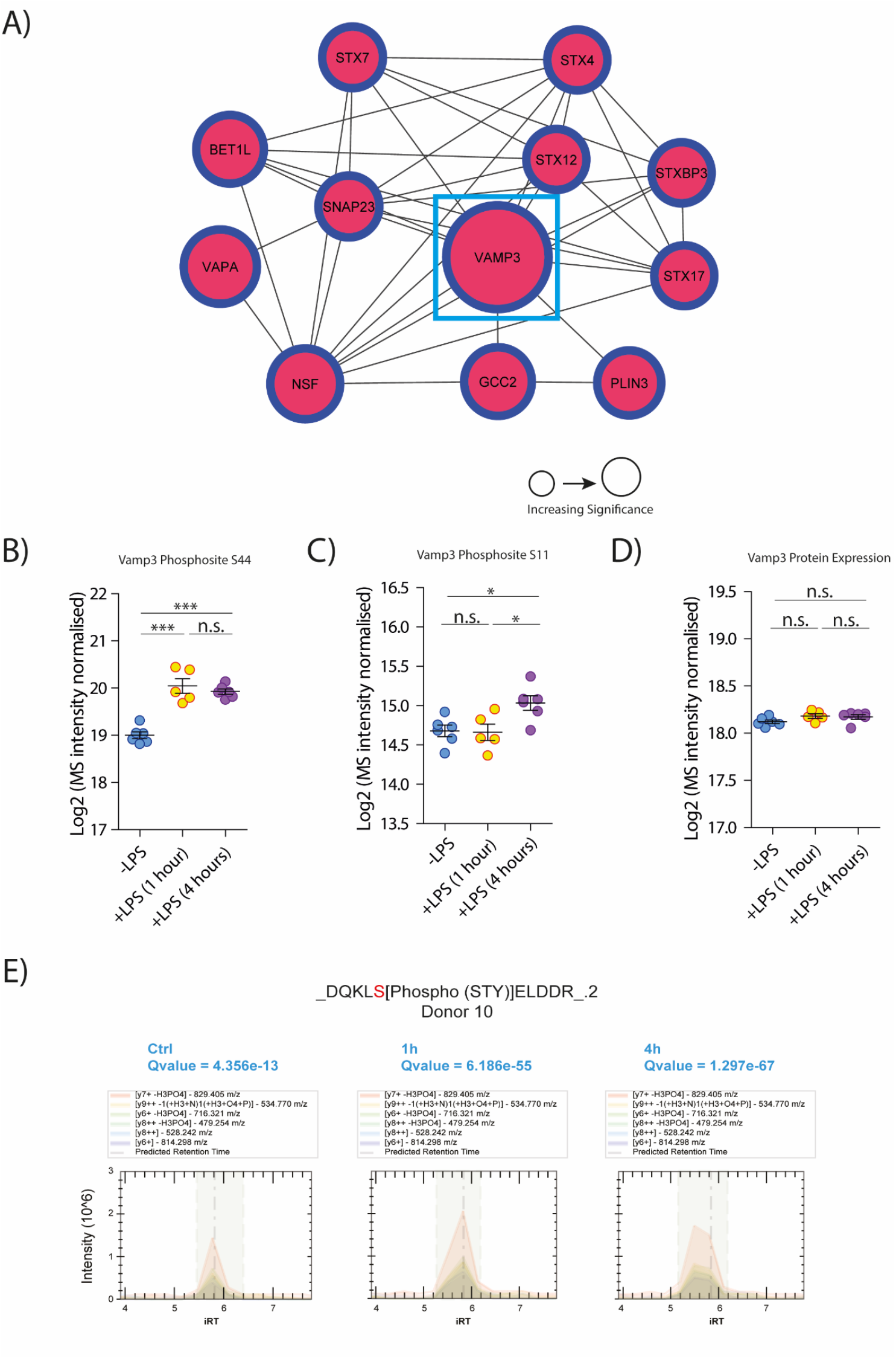
A Network of SNARE and SNARE Associated Proteins are Phosphoregulated in Activated Dendritic Cells. A). Interaction network of SNARE-related proteins found to undergo significant phosphorylation/dephosphorylation events in response to LPS stimulation (at 1 hour and/or 4 hours). Node size indicates relative significance, as calculated by 1/Q value in the original mass spectrometry dataset. B) Scatterplot of the normalized MS signal, after Log2 transformation, for the phosphorylated serine 44 of VAMP3. C) Scatterplot of the normalized MS signal, after Log2 transformation, for the phosphorylated serine 11 of VAMP3. D) Scatterplot of the normalized MS signal, after Log2 transformation, for the protein VAMP3. E) Extracted MS2 ion chromatogram for the VAMP3 serine 44 phosphopeptide. Each line represents a fragment ion. iRT = independent retention time (in minutes). ANOVA/Tukey multiple comparison tests. *, P<0.05, ***, P < 0.001; n.s., not significant.

### VAMP3 Phosphorylation Drives SNARE Complex Formation

Having established that VAMP3 is consistently phosphorylated in response to LPS stimulus at both 1 and 4 hours, we looked to establish a functional role for that phosphorylation. Previous work has shown that phosphorylation can alter the specificity of R-SNAREs to complex with Q-SNAREs (Warner, Mahajan and van den Bogaart, 2022). We therefore predicted that phosphorylation of VAMP3 drives complexing with STX4 as we previously demonstrated that VAMP3 shows enhanced complexing with STX4 in dendritic cells in response to LPS stimulation (Verboogen, Mancha, ter Beest and van den Bogaart, 2017). To test this hypothesis, we utilised Förster resonance energy transfer fluorescence lifetime imaging microscopy (FRET-FLIM). As we previously described (Linders et al., 2021; Verboogen et al., 2017), this technique can visualize SNARE complexes in living cells with organellar resolution. We used FRET-FLIM to determine if this phosphorylation would alter the fusogenic activity of VAMP3 with STX4. Specifically, STX4 and VAMP3 were C-terminally fused with mCitrine or mCherry respectively (Fig 3A). Following the complexing of VAMP3 with STX4, the C-terminal fluorophores come into close proximity, enable FRET between the donor fluorophore (mCitrine) and the acceptor fluorophore (mCherry) (Fig. 3A). This can be observed via a decrease in the fluorescent lifetime of mCitrine. This lifetime is an intrinsic property of fluorophores, and is unaffected by variables such as the local concentration of donor/acceptor fluorosphores (Linders et al., 2021). Surprisingly, we were unable to overexpress the human variant of VAMP3 in human dendritic cells, so we opted to overexpress mouse variants of fluorescently labelled VAMP3 and STX4.

**Figure 3.**
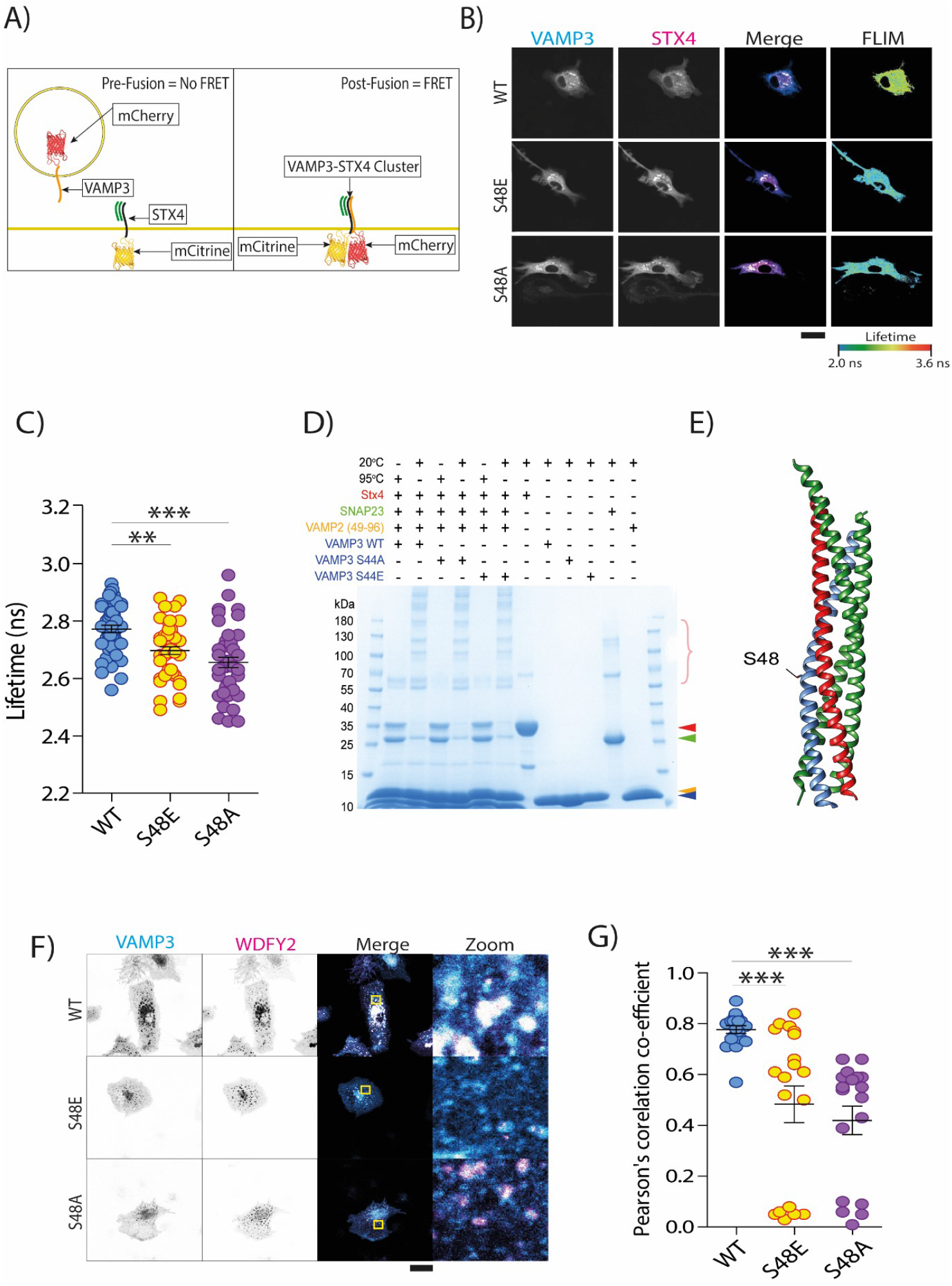
VAMP3 Phosphorylation Reprograms the VAMP3 Interactome. A) Schematic of FRET-FLIM protocol that enables the monitoring of SNARE complexing. B) Representative confocal micrographs and FLIM images of dendritic cells expressing mCitrine-STX4 and either mCherry-VAMP3 (WT), mCherry-VAMP3 (S48E) or mCherry-VAMP3 (S48A). C) Quantification of mCitrine-STX4 lifetime in dendritic cells expressing different VAMP3 mutants indicates that both the phosphomimetic (S48E) and phosphodead (S48A) VAMP3 mutants complex more efficiently with STX4, compared to VAMP3 (WT) (minimum of 43 measurements over 4 donors). D) In vitro SNARE complex formation of purified cytosolic fragments of human VAMP3 WT, S44A and S44E (blue) with a preformed acceptor complex of SNAP23 (green), cytosolic fragment of STX4 (red arrowhead) and VAMP2(49-96) (orange) analysed by SDS-PAGE and Coomassie staining. Ternary SNARE complexes (multiple bands above ∼55 kDa; pink accolade) are SDS resistant at 20°C but disassemble at 95°C. E) Crystal structure of SNARE complex with rat VAMP3(Lyubimov et al., 2016) with S48 indicated. F) Example confocal micrographs of dendritic cells overexpressing GFP-WDFY2 and VAMP3 variants. G) Pearson’s correlation coefficient of WDFY2 with VAMP3 WT or VAMP3 phosphomimetic or VAMP3 phosphodead reveals that the interaction of VAMP3 with WDFY2 is dependent on the OH group of serine 48 (minimum of 18 measurements over 2 donors). Scale bars indicate 20 microns. ANOVA/Tukey multiple comparison tests. **, P < 0.01; ***, P < 0.001; n.s., not significant.

Analysing the lifetime of mCitrine-STX4, in dendritic cells overexpressing mCherry-VAMP3 wild-type (WT), S48E or S48A (the mouse equivalent of human VAMP3 serine 44) revealed that both the phosphomimetic and phosphodead variants are more effective at complexing with STX4 at the plasma membrane, relative to the wild-type variant (Fig 3B and 3C). Unexpectedly, this fusion did not appear polarised, however this may be an artefact of protein overexpression. Thus, it seems unlikely that the phosphate group is driving the biophysical process of SNARE complexing. Rather it seems that disruption of the OH group on the serine is sufficient to enhance the complexing of VAMP3 with STX4. Therefore, we hypothesised that this OH group drives an interaction between VAMP3 and an inhibitory chaperone that blocks translocation of the associate vesicle to the plasma membrane, but does not impact complexing of VAMP3 with STX4. Consistent with this, recombinantly purified cytosolic fragments of human VAMP3 (WT, S44E and S44A) showed equal potential for complexing *in vitro* with a preformed acceptor complex of SNAP23 and the cytosolic fragment of STX4 (Fig. 3D). Here, we took advantage of the fact that SNARE complex formation is greatly accelerated when a 1:1 acceptor complex of STX4 and SNAP23 is stabilized with the C-terminal half of the SNARE motif of VAMP2 (residues 49-96) (Pobbati, Stein and Fasshauer, 2006). Moreover, in the crystal structure of the SNARE complex of rat VAMP3 with SNAP25 and STX1A (Lyubimov *et al*., 2016), S48 of rat VAMP3 (homologous to S44 of human VAMP3) points away from the SNARE bundle, supporting that this residue does not directly affect SNARE complex formation (Fig 3E).

Given that serine 44/48 does not directly affect SNARE complex formation *in vitro*, yet its loss drives complexing *in vivo*, we hypothesised that the OH group drives an interaction between VAMP3 and the chaperone WDFY2. WDFY2 has previously been shown to restrain VAMP3+ vesicle dynamics, through binding to VAMP3 (Sneeggen et al., 2019). To test this hypothesis, we co-expressed WDFY2 with either VAMP3 (WT), VAMP3 (S48E), or VAMP3 (S48A) in dendritic cells, and imaged them using confocal microscopy (Fig 3F). Whilst VAMP3 WT consistently showed strong co-localisation with WDFY2, expression of the phosphodead or phosphomimetic mutants largely led to lower WDFY2 expression (suggesting that WDFY2 is degraded in cells expressing these mutants), making co-localisation analysis impossible for most donors. Moreover, Pearson correlation analysis, which is insensitive to absolute expression levels, revealed a significant decrease in co-localisation of VAMP3 S48E or S48A with WDFY2 compared to wildtype VAMP3 (Fig 3F and 3G). These data indicate that loss of the OH-group upon phosphorylation results in dissociation of WDFY2.

In summary, these results suggest that in inactive dendritic cells, VAMP3 forms a complex with WDFY2 on VAMP3+ endosomes, restricting the traffic of the associated cargo (Fig 4A). LPS-induced activation of the dendritic cell leads to the phosphorylation of VAMP3 at serine 44 (or 48 in mice) in dendritic cells, priming the cell for the eventual synthesis of IL-6 (Fig 4B). This in turn de-stabilises the interaction of VAMP3 with WDFY2 enabling VAMP3 to complex with STX4 at the plasma membrane. Once synthesised, IL-6+ vesicles are evenly distributed throughout the cell. IL-6 is eventually shuttled to VAMP3+ recycling endosomal vesicles, which facilitate the polarised secretion of IL-6 at the plasma membrane (Fig 4C).

**Figure 4.**
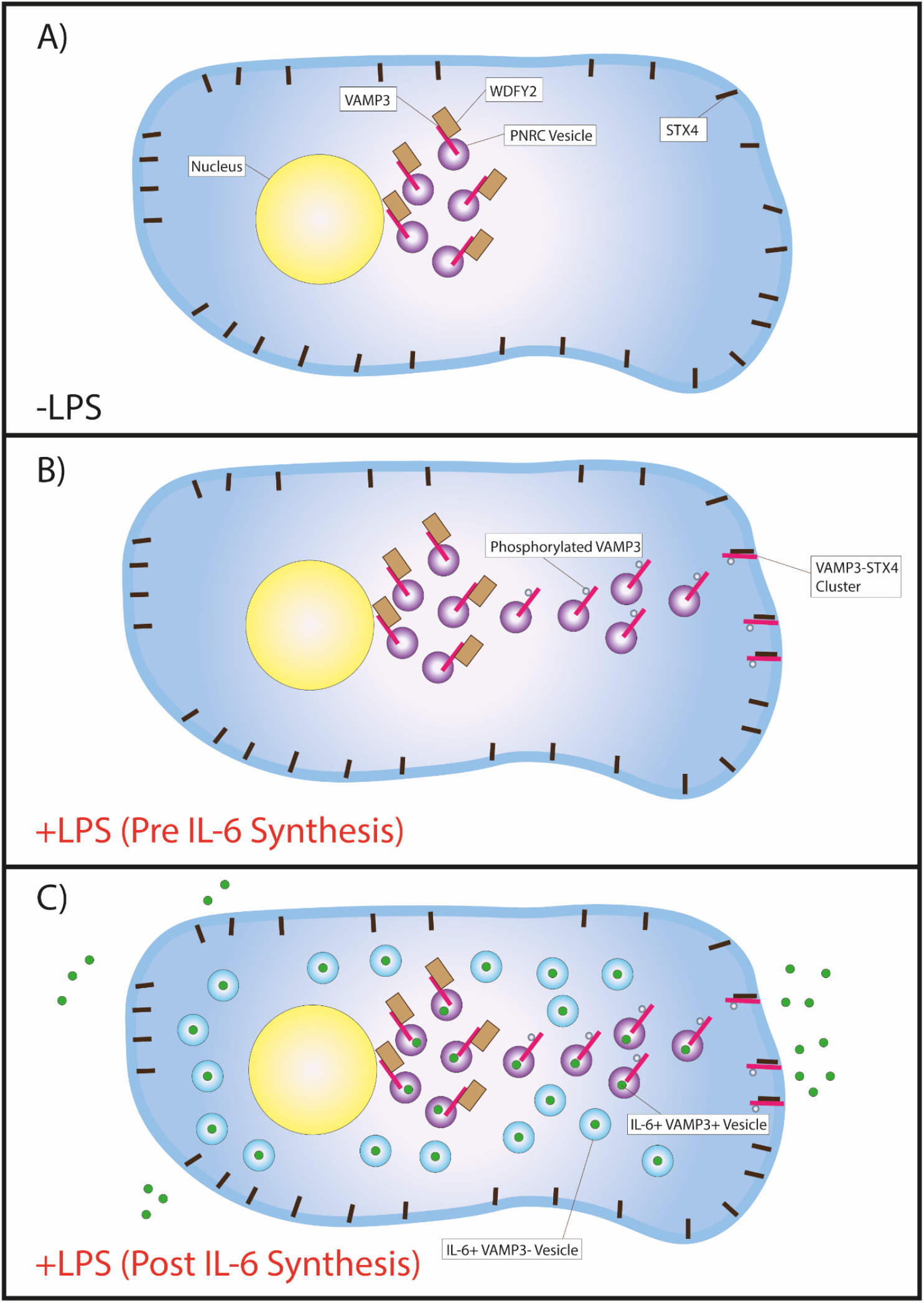
Model figure. A) Inactive dendritic cells restrict VAMP3+ perinuclear recycling compartment (PNRC) vesicle movement and/or complexing with STX4 through the formation of a WDFY2-VAMP3 complex. This interaction is dependent on the OH group of VAMP3 serine 44 (48 in mice). B) Exposure of dendritic cells to LPS primes the cell for IL-6 secretion. This occurs in part through the phosphorylation of VAMP3 at serine 44, disrupting the VAMP3-WDFY2 complex. VAMP3+ vesicles can then move to the plasma membrane, enabling VAMP3 to complex with STX4. C) Newly synthesised IL-6 is trafficked through the biosynthetic trafficking pathway to the PNRC, after which it is secreted in a VAMP-3 dependent manner. The appearance of IL-6+ VAMP3 – vesicles obscures the polarised delivery of IL-6+ VAMP3+ vesicles to the plasma membrane. This largely (but not exclusively) polarised secretion may enable myeloid cells to couple IL-6 secretion to local tissue architecture, or the polarised delivery of IL-6 to cognate T-cells.

## Discussion

Myeloid cell activation has long been understood in broad genetic terms. Elegant genetic and biochemical studies have identified key cell-surface receptors and transcription factors that trigger activation and numerous studies have de-lineated how cell-surface receptors signal to activate these transcription factors(Kawasaki & Kawai, 2014)(Tanaka, Narazaki, Masuda and Kishimoto, 2016). However, the complex molecular choreography that enables myeloid cells to perform downstream cytokine production is poorly understood. As “omics” technologies continue to develop, more of these molecular events will likely be identified. Here, we have described the phosphorylation events regulating SNARE-controlled membrane traffic in response to dendritic cell activation. Surprisingly, many alterations to trafficking machinery occur very rapidly after pathogenic stimulation; within an hour in some cases. This is particularly unusual, as this is faster than the cell can express many inflammation-associated proteins, such as cytokines (Revelo et al., 2019). This would suggest that myeloid cells prime themselves for cytokine secretion in advance of initial cytokine synthesis. This may also enable dendritic cells and macrophages to re-distribute key pre-synthesised membrane proteins in order to initiate pathogen killing. For instance, NOX2 must be continually trafficked to phagosomes in order to maintain sufficient levels of reactive oxygen species (ROS) within the phagosomal lumen (Dingjan et al., 2017).

The SNARE protein VAMP3 has long been known to present a polarised distribution within myeloid cells(Veale et al., 2011). However, the distribution and secretion of IL-6 within myeloid cells has been thought to be unpolarised (Manderson et al., 2007). This presents a paradox, as IL-6 secretion is dependent on VAMP3, which typically drives polarised cargo transport (Bajno et al., 2000; Fields et al., 2007; Verboogen et al., 2019). Here, we have resolved this paradox using TIRF microscopy. We have shown that whilst the distribution of IL-6 is unpolarised within the dendritic cell, the secretory events are polarised. This may enable dendritic cells to couple secretion to the architecture of their local environment, and may enhance their ability to activate T-cells.

SNARE phosphorylation is a long-recognised means by which cells can control and fine-tune membrane trafficking events (Warner et al., 2022). This includes immune cells, as for example phosphorylation of VAMP8 in mast cells has been proposed to act as a break on VAMP8 mediated vesicle fusion with the plasma membrane(Malmersjö et al., 2016). Furthermore, phosphorylation of SNAP23 in dendritic cells can enhance antigen cross-presentation by trafficking MHC class I molecules to the antigen-containing compartment (Nair-Gupta et al., 2014). Here we have demonstrated that phosphorylation of VAMP3 at serine 44 occurs in response to LPS stimulation in human dendritic cells and this phosphorylation drives fusion of VAMP3 positive vesicles with STX4 at the plasma membrane. Given that this phosphorylation has been observed in 19 previously published proteomic datasets (PhosphoSite Plus -Hornbeck et al., 2015), it seems likely that this represents a more general mechanism to control VAMP3 + vesicle traffic. Furthermore, this site is conserved in VAMP2, suggesting this mechanism may also exist in neurons and neuroendocrine cells, in which VAMP2 mediates neurotransmitter release (Hong, 2005)(Jahn and Scheller, 2006). Indeed WDFY2 has previously been shown to regulate VAMP2(Fritzius *et al*., 2007).

Finally, given the dual physiological and pathological roles of IL-6 in immune responses and cytokine storms respectively, it is critical that we find new ways to carefully control cytokine production in patients(Chaudhry et al., 2013). Therefore, future work should look to identify the kinase(s) that phosphorylate VAMP3 at serine 44 in response to inflammatory activation, as this may (potentially) enable IL-6 secretion to be blocked post IL-6 synthesis. Such a block in secretion could be advantageous, as it may allow for the fine-tuning of extracellular IL-6 levels in a wide variety of pathologies.

## Methods and Materials

### Plasmids

For mammalian expression, pmCitrine-N1-STX4 (Addgene, catalog number 92422) and pmCherry-N1-VAMP3 (Addgene, 92423) were generated for a previous study(Verboogen et al., 2017). pmCherry-N1-VAMP3 S48E pmCherry-N1-VAMP3 S48A and pEGFP-N1-WDFY2 (human) were generated as synthetic constructs (Genscript).

The SNAP23 and VAMP2(49-96) constructs for bacterial expression have been described previously(Antonin *et al*., 2002) (Pobbati, Stein and Fasshauer, 2006). Human VAMP3 WT, S44A and S44A (lacking the transmembrane domain, residues 1-77) and human STX4 (lacking the transmembrane domain; residues 1-271 with 272C) were generated as synthetic genes (Genscript) with codon optimization for human expression, and cloned into the NdeI/XhoI site of pET28a(+).

### Antibodies

The following primary antibodies were used in this study: rabbit polyclonal anti-VAMP3 (Abcam, Cambridge, #5789), rabbit polyclonal anti-WDFY2 (Invitrogen, #PA5-98751). The following secondary stainings were used in this study: Alexa fluor 488 Phalloidin (Invitrogen, #A12379) (1/200), Goat anti-rabbit Alex fluor-647 (Invitrogen, #A27040) (1/200).

### Cells

Monocyte-derived dendritic cells were obtained by differentiating human peripheral blood CD14+ monocytes with interleukin (IL)-4 (300 µg/ml) and granulocyte-macrophage colony-stimulating factor (GM-CSF; 450 µg/ml) for 6 days in RPMI supplemented with 10% serum, antibiotics (100 μg ml^−1^ penicillin, 100 µg ml^−1^ streptomycin and 0.25 µg ml^−1^ amphotericin B, Gibco), and 2 mM glutamine. Monocytes were isolated from the blood of healthy donors (informed consent and consent to publish obtained, approved by the ethical committee of Dutch blood bank Sanquin) as previously described (de Vries et al., 2002). LPS (O111:B4, Sigma Aldrich 32160405) stimulation was carried out overnight unless otherwise stated.

### Confocal Microscopy

Cells were seeded onto glass coverslips for at least 12 hours before fixation. Cells were fixed in 4% PFA for 15 minutes at room temperature, or ice-cold methanol for 10 minutes at -20°C (for lamin-A/C immunolabeling). Cells fixed in PFA were permeabilized for 5 minutes in a 0.1% (v/v) Triton-X100 solution. Cells were blocked in a 20 mM glycine 3% BSA PBS-based solution for 1 hour before antibody staining.

Transient transfection of dendritic cells was achieved using the Neon-transfection system (Thermo Scientific). 1.2 million cells were washed with PBS and suspended in 115 µL of buffer R with 5 µg of DNA. Cells were pulsed twice for 40 ms at 1000 V. Cells were then transferred to phenol red-free RPMI with 20% serum for at least 4 hours before imaging. Live cell imaging was performed in OptiKlear solution (Abcam ab275928).

Images were collected with a Zeiss LSM 800 microscope equipped with a Plan-Apochromat (63x/1.4) oil DIC M27 (FWD=0,19 mm) objective (Zeiss). Images were acquired using the ZEN software (version 2.3). DAPI was excited by a 405 laser, and Alexa Fluor 488 phalloidin was excited by a 488 laser. For Z-series, a slice interval of 0.31 µm was used. Airyscan microscopy was performed with a Zeiss LSM 800 airyscan microscope. A Z-interval of 0.17 µm was used. Images were acquired using the ZEN software (version 2.3). Images were subject to airyscan processing following acquisition.

### TIRF microscopy and analysis

Total internal reflection fluorescence (TIRF) was performed on Olympus IX51 inverted microscope equipped with a 60x/1.49 oil immersion objective cell TIRF illuminator (Olympus). Excitation was done with a 488 nm laser. Fluorescence emission was collected with an appropriate dichroic mirror and Photometrics Prime Express sCMOS camera.

Images were recorded at 37°C as time-lapse movies of 1000 frames over an interval of 100 ms. The number of bursts/ flashes were analysed from time-lapse movies of moDCs expressing IL-6-GFP using the ImageJ plugin xySpark(Steele & Steele, 2014).

The data from xySpark (frame number, number of sparks, time) were imported into Graphpad Prism (v 8.4.3) to plot graphs and perform statistical analysis. We performed calibration to convert the cell surface area obtained in pixels using ImageJ, to obtain the cell area in µm. The results are represented as the number of sparks across different conditions and as bursts s^-1^ µm^-2^.

### FRET-FLIM

Live imaging was performed on a Leica SP8 SMD system at 37 °C, equipped with an HC PL APO CS2 63x/1.20 Water objective. Fluorophores were excited with a pulsed white-light laser, operating at 80 MHz. mCitrine was excited at 514 nm, two separate HyD detectors were used to collect photons, set at 521–565 nm and 613–668 nm, respectively. Photons were collected for 1 min and lifetime histograms of the donor fluorophore were fitted with monoexponential decay functions convoluted with the microscope instrument response function in Leica LAS X. Data analysis was performed with the FLIMfit software tool (version 5.1.1) developed at Imperial College London. All data was fitted with exponential decay functions convoluted with an instrument response function (IRF). The lifetime of mCitrine-STX4 was then analysed at the cell periphery (approximating the plasma membrane).

### *In vitro* complex formation

SNAREs were expressed in BL21 (DE3), induced at OD_600_ = ∼0.5 with Isopropyl B-D-1-thiogalactopyranoside (IPTG) in a final concentration of 0.5 mM for 6 h. Cell pellets were incubated for 30 min at room temperature (RT) at 4 ml/per gram with extraction buffer: 20 mM HEPES pH 7.4, 500 mM NaCI, 8 mM imidazole, 1 mM Tris(2-carboxyethyl)phosphine hydrochloride (TCEP), 1 mM MgCl, 1 mM phenylmethanesulfonyl fluoride (PMS), DNAse (2 μg/ml) and lysozyme (0.2 mg/ml)). The same volume of extraction buffer as used before with an additional 10% of sodium cholate hydrate (3a,7a, 12a-Trihydroxy-5B-cholan-24-oic acid sodium salt) was added. After another incubation step of 15 min at RT, 4M urea was added and the reaction was further incubated for 30 min at RT. The cells were lysed via sonication (MSE, Soniprep 150 Plus) with 10 cycles of 15 seconds sonication (Amplitude: 6-8, 30 seconds off and on ice). After taking a sample, cell debris was removed via centrifugation (Thermo Scientific, Sorval Lynx 40001) at 27,000 x g at 4 °C for 30 min. After taking a sample from the cell-free extract (CFE) the chromatography column (BioRad, Poly-Prep Chromatography Column) was prepared by adding 1 ml of resin (His-select nickel affinity gel), washing the column twice with 5 ml of ice-cold MilliQ and equilibrating it with 5 ml of wash buffer: 20 mM HEPES pH 7.4, 500 mM NaCI, 20 mM imidazole, 1 mM TCEP, 1% 3-[(3-Cholamidopropyl)dimethylammonio]-1-propanesulfonate hydrate (CHAPS).

CFE was added to the column and mixed well with the resin. After incubating for 2 h on a roller at 4 C°, the column was opened and a sample from the flow-through was collected. The column was washed 5 times with 500 µl of wash buffer and then 5 times with 500 µl elution buffer (20 mM HEPES pH 7.4, 500 mM NaCI, 400 mM imidazole, 1 mM TCEP, 1% CHAPS). Samples were collected and the concentration was measured by absorption at 280 nm. 50 µl of thrombin (5 mg/ml in 50% glycerol = 1 U/ml) was added to the elution with the highest concentration, in order to cleave the His-tag from the SNARE protein. The reaction was performed in dialysis cups (Thermo Scientific, Slide-A-Lyzer MINI Dialysis Device, 2K MWCO, 0.1 ml) with dialysis buffer (20 mM HEPES pH 7.4, 100 mM NaCl, 0.2 mM TCEP and 1% CHAPS) overnight at 4 °C in a beaker while stirring.

For further purification of VAMP3 mutants, cation exchange chromatography (BioRad, NGC QuestTM 10 Chromatography System #7880001) was performed. As a column, the SOURCE 15S 4.6/100 EP (cation) was used. Prior to loading, the column was equilibrated with AKTA-A buffer (20 mM HEPES pH 7.4, 1 M NaCl, 0.2 mM TCEP and 1% CHAPS). For elution, a gradient volume of 15 column volumes to a Sodium chloride concentration of 0.5 M AKTA-B buffer (AKTA-A with 1 M NaCl) was used. This segment was followed by stepping up the gradient to a salt concentration of 1 M (100% AKTA-B) and then holding it at 1 M for 3 ml before re-equilibration the column with 3 ml of AKTA-A. Before storing the fractions at -20 °C, glycerol was added in a final concentration of 10% as a lyoprotectant.

For further purification of the STX4 complex with SNAP23 and VAMP2(49-96), 100 mM of each protein was mixed. The mixture was incubated overnight at RT. The complex was then purified using size exclusion chromatography (BioRad, NGC QuestTM 10 Chromatography System #7880001). Prior to loading, the column (ENrich SEC 70 10 × 300 mm column) was equilibrated with dialysis buffer (20 mM HEPES pH 7.4, 100 mM NaCl, 0.2 mM TCEP and 1% CHAPS). The protein concentration of the elution fractions was determined by BCA Assay (Pierce BCA Protein Assay Kit).

For SNARE complex formation, 10 mM of the STX4 complex with SNAP23 and VAMP2{49-96) was mixed with VAMP3 and incubated at room temperature overnight. Further analysis was done by SDS-PAGE and Coomassie staining. Samples were mixed with 2x Laemmli buffer and heated for 10 min at 95 °C or not heated. BioRad precast protein gels (4-20% Mini-PROTEAN TGX Precast Protein Gels,15-well, 15L #4561096) were used for the SDS-PAGE.

